# Impaired carotid body hypoxic sensing in mice deficient in olfactory receptor 78

**DOI:** 10.1101/757120

**Authors:** Ying-Jie Peng, Anna Gridina, Jayasri Nanduri, Aaron P. Fox, Nanduri R. Prabhakar

## Abstract

Carotid bodies are the sensory organs for detecting hypoxemia (decreased arterial blood oxygen levels) and ensuing chemo reflex is a major regulator of breathing and blood pressure. Chang et al (2015) proposed that olfactory receptor 78 (Olfr78) plays a major role in hypoxic sensing by the carotid body. However, such a possibility was questioned by a subsequent study ((Torres-Torrelo et al. 2018). The discrepancy between the two reports prompted the present study to re-examine the role of Olfr78 in hypoxic sensing by the carotid body (CB). Studies were performed on age and gender matched Olfr78 knock out mice generated on BL6 and JAX backgrounds and corresponding wild type mice. Breathing was monitored by plethysmography in un-sedated and efferent phrenic nerve activity in anesthetized mice. Carotid body sensory nerve activity was determined *ex vivo* and [Ca^2+^]_i_ responses were monitored in isolated glomus cells, the primary O_2_ sensing cells of the carotid body. Olfr78 null mice on both BL6 and JAX backgrounds exhibited attenuated hypoxic ventilatory response, whereas breathing responses to CO_2_ were unaffected. The magnitude of hyperoxia-induced depression of breathing (Dejour’s test), which is an indirect measure of carotid body hypoxic sensing, was markedly reduced in Olfr78 mutant mice on both background strains. Furthermore, carotid body sensory nerve and glomus cell [Ca^2+^]_i_ responses to hypoxia were attenuated in BL6 and JAX Olfr78 null mice. These results suggest that Olfr78 plays an important role in hypoxic sensing by the carotid body.

## INTRODUCTION

Carotid bodies (CB) are the sensory organs for detecting decreased arterial blood O_2_ levels. Hypoxemia increases the CB sensory nerve activity, and triggers reflex stimulation of breathing and blood pressure to ensure optimal oxygen delivery to tissues (Kumar and Prabhakar 2012). The CB is comprised of two major cell types; glomus or type I cells, which are of neuronal lineage, and sustantecular or type II cells, resembling glial cells of the nervous system. Glomus cells which are in synaptic contact with the afferent nerve ending are considered the primary site of transduction of the hypoxic stimulus (Kumar and Prabhakar 2012). There is considerable debate regarding the sensory transduction in glomus cells (Prabhakar, Peng and Nanduri 2018, Kumar and Prabhakar 2012, Rakoczy and Wyatt 2017).

Olfactory receptors (OR) are G-protein coupled receptors for detecting smell sensation (Schild and Restrepo 1998). Although ORs are originally described in olfactory neurons, emerging evidence suggest that some ORs are also expressed in other tissues (Maßberg and Hatt 2018). Chang et al reported high abundance of *Olfr 78* (also known as MOL2.3 and MOR 18-2) gene expression in glomus cells of the mouse carotid body (Chang et al. 2015), a finding that was subsequently confirmed using single cell mRNA analysis in neonatal mice (Zhou et al. 2016). Olfr78 knockout mice exhibit blunted CB sensory nerve excitation, [Ca^2+^]_i_ elevation of glomus cells by hypoxia and these effects were associated with reduced hypoxic ventilatory response (HVR), which is a hallmark of the CB chemo reflex (Chang et al. 2015). However, a subsequent study reported that HVR, [Ca^2+^]_i_ and neurotransmitter secretory response of glomus cells to hypoxia were unaffected in Olfr78 knockout mice, thereby questioning the role of Olfr78 signaling in the carotid body hypoxic sensing (Torres-Torrelo et al. 2018). The discrepancy between the studies by Chang et al (2015) and Torres-Torrelo et al (2018) prompted us to re-examine the role of Olfr78 signaling in the CB response to hypoxia. Our results showed that HVR monitored either in conscious or anesthetized mice were reduced in Olfr78 knockout mice. The attenuated HVR was associated with impaired CB sensory nerve excitation in response to graded hypoxia as well as hypoxia-evoked [Ca^2+^]_i_ elevation in glomus cells. Our results suggest that Olfr78 is an important component of the CB response to hypoxia.

## METHODS

### Preparation of Animals

Experimental protocols were approved by the Institutional Animal Care and Use Committee of the University of Chicago. Experiments were performed on age-matched adult wild-type (WT) and Olfr78 null mice on C57BL/6 background (BL6, gift from Dr. J. Pluznick, Johns Hopkins University) and on 129P2/OlaHsd background (JAX, gift from Dr. A. Chang, The University of California, San Francisco, UCSF).

### Measurements of breathing

#### Whole body plethysmography

Ventilation was monitored by whole-body plethysmograph (Buxco, DSI, St. Paul, MN), and O_2_ consumption and CO_2_ production were determined by the open-circuit method in un-sedated mice as described (Peng et al. 2006). Ventilation was recorded while the mice breathed 21% or 12% O_2_-balanced N_2_. Each gas challenge was given for 5 min. O_2_ consumption and CO_2_ production were measured at the end of each 5-min challenge. For determining ventilatory response to CO_2_, baseline ventilation was determined while mice breathed 100% O_2_ followed by hypercapnic challenge with 5% CO_2_–95% O_2_–balance N_2_. Sighs, sniffs, and movement-induced changes in breathing were monitored and excluded in the analysis. All recordings were made at an ambient temperature of 25 ± 1 °C. Minute ventilation (V_*E*_ *=* Tidal volume, V_T_ x respiratory rate, RR) was calculated and normalized for body weight and expressed as ratio to O_2_ consumption (V_*E*_ / VO_2_).

#### Measurements of efferent phrenic nerve activity

Mice were anaesthetized with intraperitoneal injections of urethane (1.2g/kg). Supplemental doses (10% of the initial dose of anesthetic) were given when corneal reflexes or responses to toe pinch were observed. Animals were placed on a warm surgical board and a tracheotomy was performed through a midline neck incision. The trachea was cannulated and mice were allowed to breathe spontaneously. Core body temperature was monitored by a rectal thermistor probe and maintained at 37±1°C by a heating pad. The phrenic nerve was isolated unilaterally at the level of the C3 and C4 spinal segments, cut distally, and placed on bipolar stainless steel electrodes. Integrated efferent phrenic nerve activity was monitored as an index of central respiratory neuronal output. The electrical activity was filtered (band pass 0.3–1.0 kHz), amplified (P511K, Grass Instrument, West Warwick, RI), and passed through Paynter filters (time constant of 100 ms; CWE Inc.) to obtain a moving average signal. Data were collected and stored in the computer for further analysis (PowerLab/8P, AD Instruments Pty Ltd, Australia). Phrenic nerve bursts/ min (index of respiratory rate, RR, tidal phrenic nerve activity, and minute neuronal respiration (MNR = RR x tidal phrenic nerve activity) were analyzed. The effects of hypoxia (12% O_2_ balanced nitrogen) and hypercapnia (5% CO_2_ balanced 95% O_2_) on efferent phrenic nerve activity were determined. Gases were administered through a needle placed in the tracheal cannula and gas flow was controlled by a flow meter. For hypoxic response, baseline phrenic nerve activity was monitored while the animals breathed room air for 3 min. Subsequently, inspired gas was switched to 12% O_2_ for 3min. The duration of 3min for hypoxia was chosen because longer duration of hypoxic exposure (> 5min) in anesthetized mice leads to hypotension which confounds interpretation of results. For hypercapnic response, 3 min of 5% CO_2_ balanced 95% O_2_ was preceded by exposure to 100% O_2_ for 3 min. At the end of the experiment, mice were killed by overdose of urethane (>3.6 g/kg, i.p.).

### Hyperoxic Challenge (Dejours test)

These experiments were performed on anesthetized mice. Baseline neural respiration (efferent phrenic nerve activity) was recorded while animals breathed 21% O_2_ for 45 sec followed by 100% O_2_ for 20 sec. Respiratory variables were analyzed during 21% O_2_ breathing and during the last 15 sec of hyperoxia (the initial 5 sec was excluded because of the dead space).

### Recording of CB sensory nerve activity

Sensory nerve activity from CB *ex vivo* was recorded as previously described (Peng et al. 2006). Briefly, CBs along with the sinus nerves were harvested from anaesthetized mice, placed in a recording chamber (volume, 250μl) and superfused with warm physiological saline (35°C) at a rate of 3 ml/ minute. The composition of the medium was (mM): NaCl, 125; KCl, 5; CaCl_2_,1.8; MgSO_4_,2; NaH_2_PO_4_, 1.2; NaHCO_3_, 25; D-Glucose, 10; Sucrose, 5. The solution was bubbled with 21% O_2_/5% CO_2_. Hypoxic/hypercapnic challenges were achieved by switching the perfusate to physiological saline equilibrated with desired levels of O_2_. To facilitate recording of clearly identifiable action potentials, the sinus nerve was treated with 0.1% collagenase for 5 min. Action potentials (1–3 active units) were recorded from one of the nerve bundles with a suction electrode and stored in a computer via a data acquisition system (PowerLab/8P). ‘Single’ units were selected based on the height and duration of the individual action potentials using the spike discrimination module. Oxygen level of the superfusate medium in the chamber was continuously monitored with a platinum electrode placed next to the CB using a polarographic amplifier (Model 1900, A-M Systems, Sequim, WA).

### Primary cultures of glomus cells

Primary cultures of glomus cells were prepared as described previously (Makarenko et al. 2012). Briefly, CBs were harvested from urethane (1.2 g/kg, I.P.) anesthetized mice. Glomus cells were dissociated using a mixture of collagenase P (2 mg/ml; Roche, USA), DNase (15μg/ml; Sigma, USA), and bovine serum albumin (BSA; 3 mg/ml; Sigma, USA) at 37°C for 20 min, followed by a 15 min incubation in medium containing DNase (30μg/ml) only. Cells were plated on collagen (type VII; Sigma)-coated cover slips and maintained at 37°C in a 7% CO_2_ + 20% O_2_ incubator for overnight (∼18h). The growth medium consisted of F-12K medium (Invitrogen) supplemented with 1% fetal bovine serum, insulin-transferrin-selenium (ITS-X; Invitrogen), and 1% penicillin/streptomycin/glutamine mixture (Invitrogen).

### Measurements of [Ca ^_2+_^]_i_

Glomus cells were incubated in Hanks Balanced Salt Solution (HBSS) with 2μM fura-2-AM and 1mg/ml albumin for 30 min and then washed in a fura-2-free solution for 30 min at 37°C. The cover slip was transferred to an experimental chamber for determining the changes in [Ca^2+^]_i_. Background fluorescence at 340nm and 380 nm wavelengths was obtained from an area of the cover slip that was devoid of cells. On each cover slip, glomus cells were identified by their characteristic clustering and individual cells were imaged. Image pairs (one at 340nm and the other at 380nm wavelength, respectively) were obtained every 2s by averaging 16 frames at each wavelength. Data were continuously collected throughout the experiment. Background fluorescence was subtracted from the cell data obtained at the individual wavelengths. The image obtained at 340nm was divided by the 380nm image to obtain ratiometric image. Ratios were converted to free [Ca^2+^]_i_ using calibration curves constructed *in vitro* by adding fura-2 (50μM free acid) to solutions containing known concentrations of Ca^2+^ (0 –2000 nM). The recording chamber was continually superfused with warm solution (31°C) from gravity-fed reservoirs. The composition of the medium was the same as that employed for recording CB sensory activity described above.

#### Data analysis

The following variables were analyzed in un-anaesthetized mice: tidal volume (VT; μl); respiratory rate (RR/min); minute ventilation (V_*E*_; ml/min·g body weight); O_2_ consumption (V_*O2*_; ml/ min); CO_2_ production (VCO_2_; ml/min). In agreement with other investigators (Tankersley, Fitzgerald and Kleeberger 1994, Dauger et al. 1998, Tankersley et al. 1993), we also found that breathing is unstable in un-sedated mice due to movement, sniffing and exploratory behaviors. Therefore, 15-20 consecutive stable breaths per minute were chosen for analysis of respiratory variables during breathing room air and during 5 min of inspired O_2_ and CO_2_ challenges. V_*T*_ and V_*E*_ were normalized to the body weight of the animals. Each data point represents the average of two trials in a given animal for a given gas challenge. In anaesthetized animals, the following respiratory variables were analyzed: respiratory rate (RR; phrenic bursts per minute), amplitude of the integrated tidal phrenic nerve activity (a.u., arbitrary units) and minute neural respiration (MNR) (number of phrenic bursts per min, RR × amplitude of the integrated tidal phrenic nerve activity, a.u.). CB sensory activity (discharge from ‘single’ units) was averaged during 3 min of baseline and during the 3 min of gas challenge and expressed as impulses per second unless otherwise stated. All data are presented as Box-Whisker plots with individual data points, unless otherwise stated. Statistical significance was assessed by either Mann-Whitney Rank Sum test or two-way ANOVA with repeated measures followed by Tukey’s test. P values < 0.05 were considered significant.

## RESULTS

### Impaired hypoxic ventilatory response (HVR) in Olfr78 null mice

In the first series of experiments, breathing was monitored by plethysmography in un-sedated mice. Examples of the hypoxic ventilatory response (HVR) and summary data of V_E_/VO_2_ in both WT and Olfr78 null mice (on BL6 and JAX backgrounds) as percentage of normoxia is presented in Fig. 1 A-D. Absolute values of respiratory changes and metabolic variables 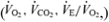 are given in Table 1A and B. Baseline breathing (21% O_2_) was comparable between WT and Olfr78 null mice (BL6 and JAX background strains). Minute ventilation presented as V_*E*_/V_*O2*,_ a measure of convective requirement, increased in response to 12% O_2_ in WT mice, which was primarily due to increased respiratory rate (Fig.1 A-D; Table 1 a-b). The magnitude of HVR was significantly attenuated in Olfr78 null mice on both BL6 and JAX backgrounds (Fig.1 A-D; Table 1 a-b).

**Table 1.**
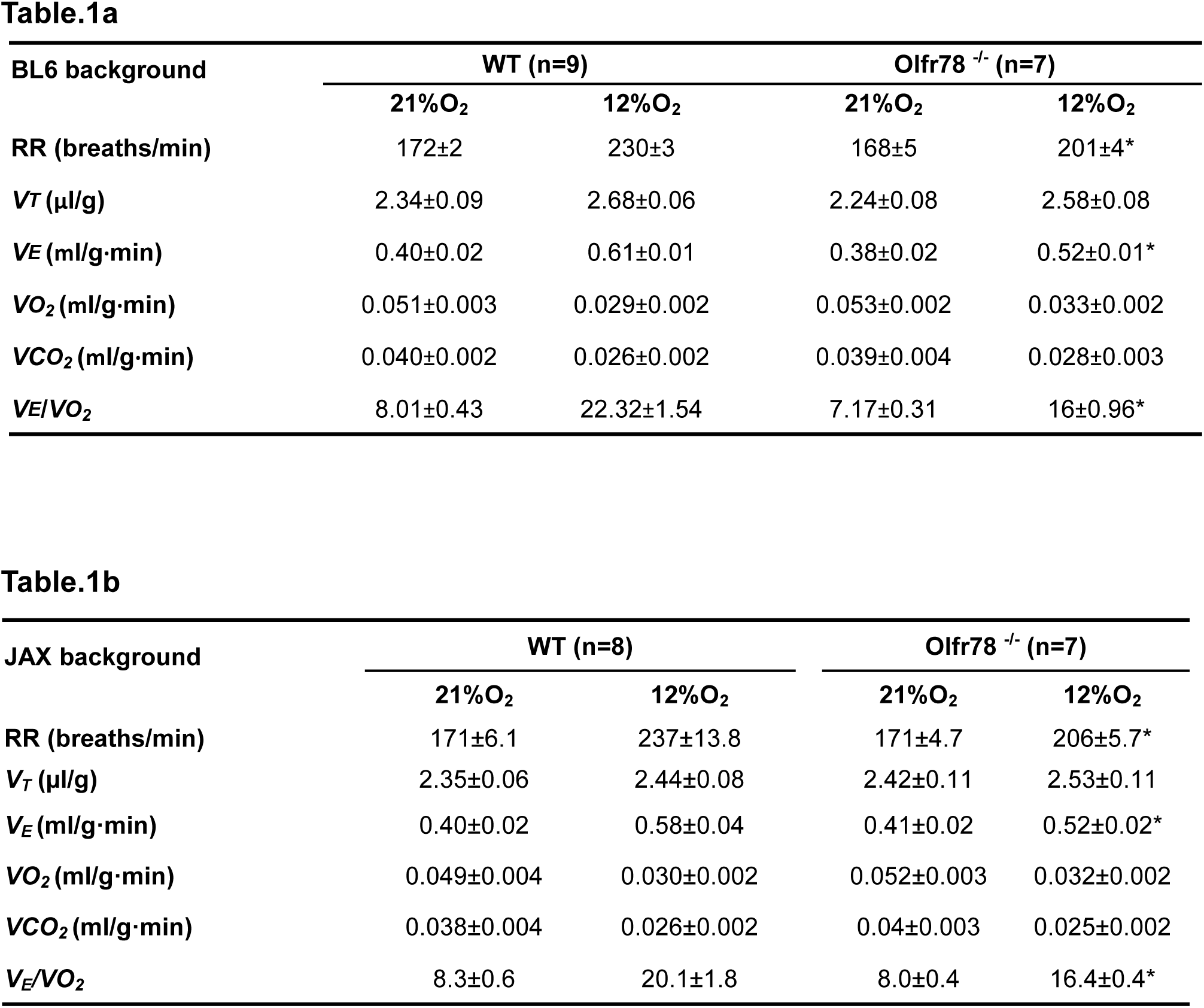
Respiratory and metabolic variables in un-sedated wild-type and Olfr78 null mice on BL6 (a) and JAX (b) backgrounds. Respiration was monitored by whole body plethysmography. Numbers in parenthesis represent number of mice. RR, respiratory rate per min, *V*_*T*_, tidal volume (µl), minute ventilation,V_E_, Oxygen consumption, V_O2_, CO_2_ production, WT vs Olfr78 Two-way ANOVA with repeated measures followed by Tukey test, * p< 0.05.

**Fig. 1.**
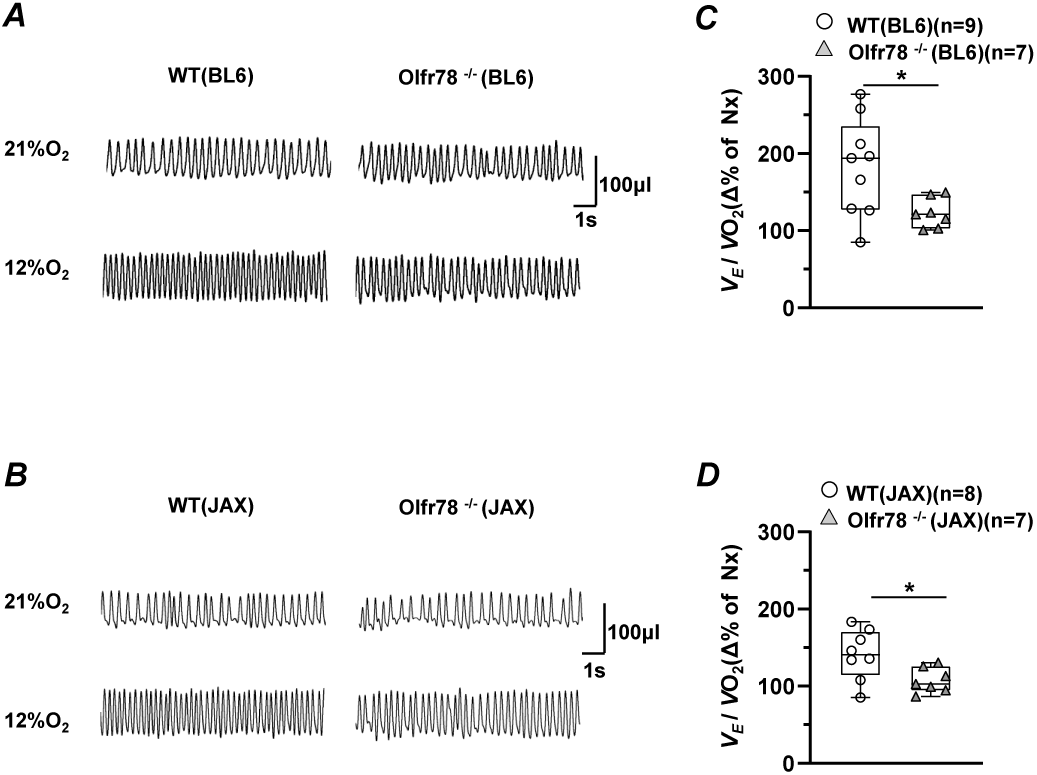
Ventilatory responses to hypoxia in wild type (WT) and Olfr 78 ^−/−^ mice on BL6 and JAX backgrounds. Ventilation was measured in un-sedated mice by whole body plethysmography under normoxia (21% O_2_) and hypoxia (12% O_2_) along with O_2_ consumption (V_O2_). Hypoxia was given for 5 min. Representative tracings of breathing are shown in ***A*** and ***B.*** Summary data of minute ventilation normalized to body weight and oxygen consumption (V_E_/V_O2_), later representing measure of convective requirement are shown as Box-Whisker plots with individual data points in ***C*** and ***D***. In panels ***C*** and ***D***, parenthesis represents number of mice in each group. WT vs Olfr78 null mice, Mann-Whitney Rank Sum test, * P < 0.05.

In the second series of experiments, efferent phrenic nerve activity was monitored as an index of respiratory neuronal output in anesthetized mice. Examples of neural breathing responses to hypoxia (12% O_2_) in WT and Olfr78 null mice on BL6 and JAX background and summary data of minute neural respiration during hypoxia are presented in Figure 2 A-D. WT mice showed increased minute neural respiration during hypoxia, which was due to increased phrenic burst frequency (RR) as well as tidal phrenic amplitude and these responses were markedly attenuated in Olfr78 null mice (Fig.2 A-D).

**Fig.2.**
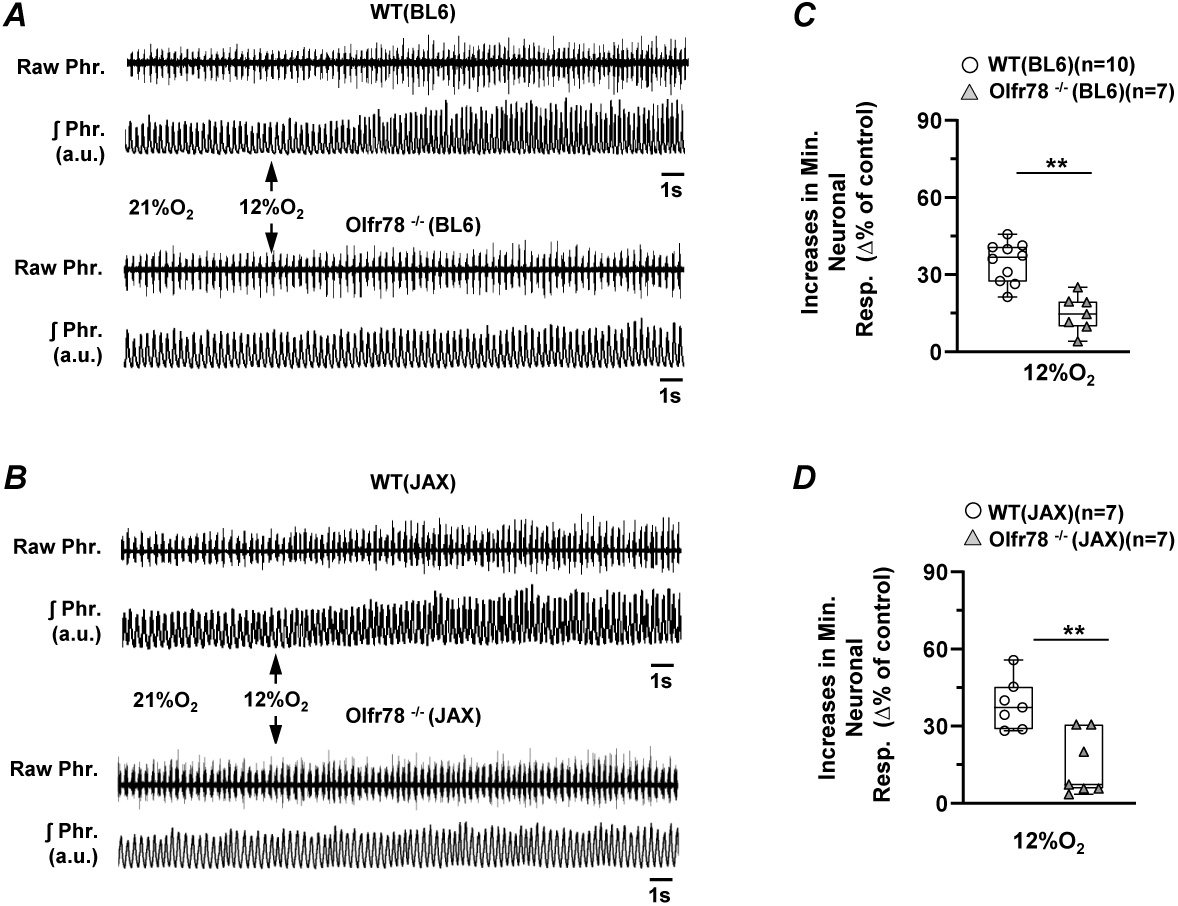
***A***-***B***. Examples of efferent phrenic nerve activity during 21 and 12% O_2_ (at arrow) in anesthetized, spontaneously breathing wild-type (WT) (*Upper panels*) and Olfr78 null mice (*Lower panels*) on BL6 and JAX backgrounds. Raw Phr., action potentials of the phrenic nerve activity. ∫ Phr., integrated phrenic nerve activity (a.u. arbitrary units). ***C-D***. Box-Whisker plots with individual data points showing data of minute neural respiration (number of phrenic bursts/min x tidal phrenic activity) presented as percent of baseline activity during 21% O_2_ breathing. Parenthesis represents number of mice in each group. WT vs Olfr78 null mice, Mann-Whitney Rank Sum test, **, P < 0.01.

In contrast, WT and Olfr78 null mice (BL6 and JAX background) responded to hypercapnia (5% CO_2_) with comparable increases in breathing measured either by plethysmography (Fig. 3 A-D) or by efferent phrenic nerve activity (Fig. 4 A-D).

**Fig.3.**
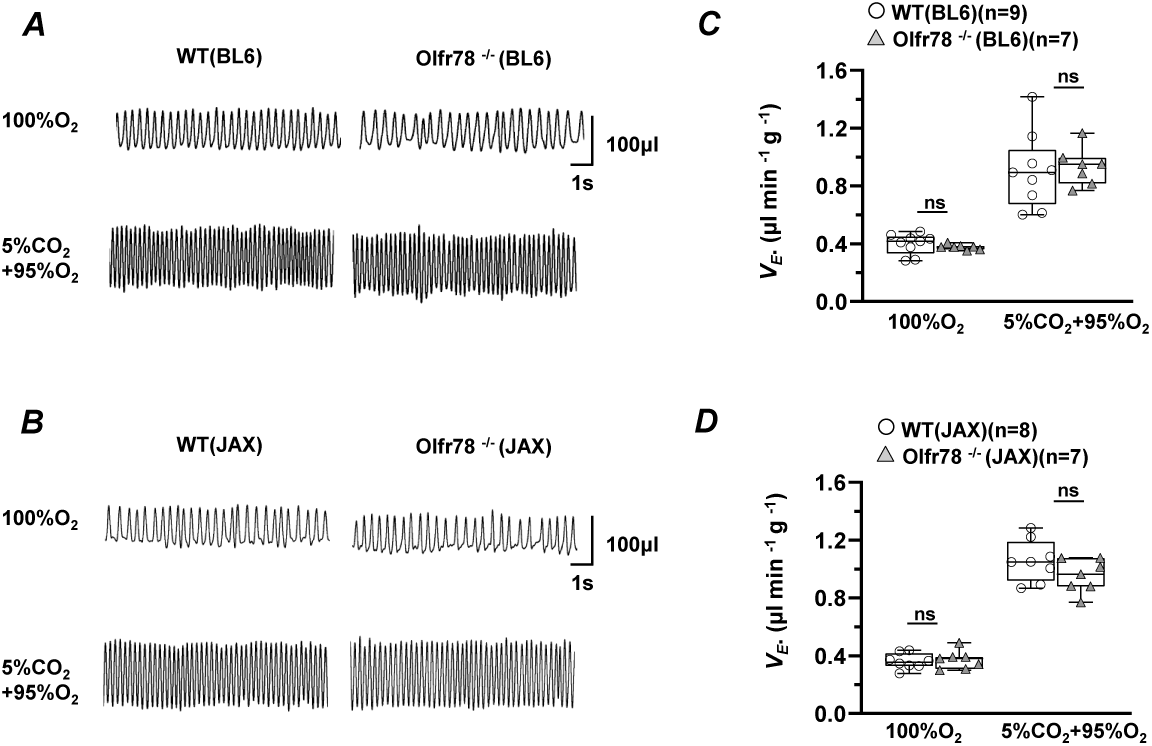
Ventilatory responses to hypercapnia (5% CO_2_) in wild type (WT) and Olfr 78 ^−/−^ mice on BL6 and JAX backgrounds. Ventilation was measured in un-sedated mice by whole body plethysmography under hyperoxia (100% O_2_), and hypercapnia (5%CO_2_ +95% O_2_). Hypercapnia challenge was given for 5 min. Representative tracings of breathing are shown in ***A*** and ***B***. Summary data of minute ventilation (V_E_) normalized to body weight in response to hypercapnia are shown in ***C*** and ***D*** as Box-Whisker plots with individual data points. Parenthesis represents number of mice in each group. WT vs Olfr78 null mice, Two-way ANOVA with repeated measures followed by Tukey test, n.s. not significant, P > 0.05 as compared to WT.

**Fig. 4.**
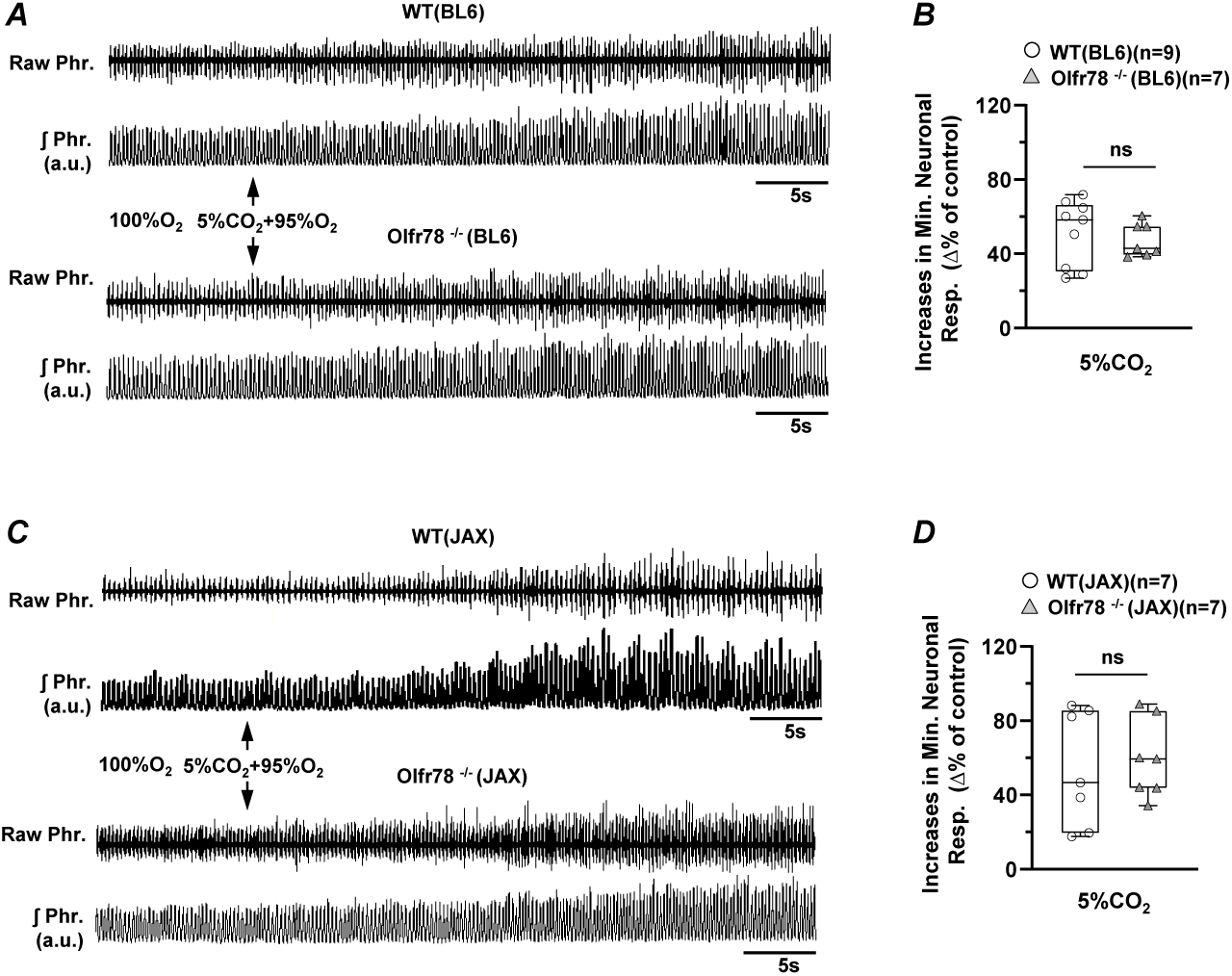
***A***-***B***. Efferent phrenic nerve activity during 100 % O_2_ and 5% CO_2_+95% O_2_ breathing (at arrow) in anesthetized, spontaneously breathing wild-type (WT) (*Upper panels*) and Olfr78 null mice (*Lower panels*) on BL6 and JAX backgrounds, respectively. Raw Phr., action potentials of the phrenic nerve activity. ∫ Phr., integrated phrenic nerve activity (a.u. arbitrary units). ***C-D***. Box-Whisker plots with individual data points showing data of minute neural respiration (number of phrenic bursta/min x amplitude of tidal phrenic activity in a.u.) presented as percent of baseline breathing during 100% O_2_ breathing. Parenthesis represents number of mice in each group. WT vs Olfr78 null mice, Mann-Whitney Rank Sum test, n.s. not significant, P > 0.05.

### Assessment of carotid body function in Olfr78 null mice

The magnitude of the transient depression of breathing in response to a brief hyperoxic exposure is considered as an index of peripheral chemoreceptor, especially the CB sensitivity to O_2_ (Dejours 1962). Phrenic nerve activity was monitored in anesthetized mice for 45sec while breathing room air and then for 20 sec after a hyperoxic challenge (100% O_2_). As compared with Olfr78 null mice (BL6 and JAX backgrounds), wild-type mice manifested a significantly greater depression of minute neural respiration in response to hyperoxia (Fig. 5 A-D).

**Fig. 5.**
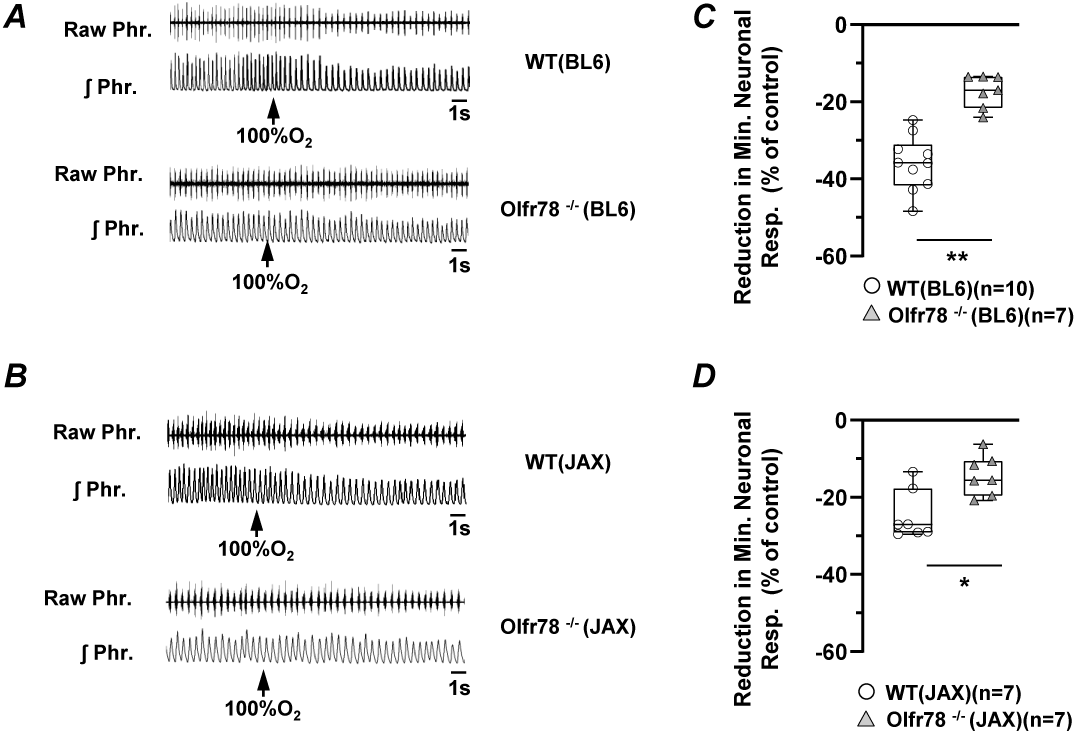
***A-B***. Efferent phrenic nerve responses to brief (20 s) exposure to 100%O_2_ (at arrow) in anesthetized, spontaneously breathing wild-type (WT) (*Upper panels*) and Olfr78 null mice on BL6 and JAX background (*Lower panels*). Raw Phr., action potentials of the phrenic nerve activity. ∫Phr., integrated efferent phrenic nerve activity. ***C-D***. Box-Whisker plots with individual data points showing data of hyperoxia-induced reduction in minute neural respiration (number of phrenic bursts/min x amplitude of tidal phrenic nerve activity in arbitrary units) presented as percent of baseline breathing in room air. Number of mice are presented in parenthesis. Wild type (WT) vs Olfr78 null mice, Mann-Whitney Rank Sum test, **, P < 0.01, * P<0.05.

To further establish the effects of hypoxia, CBs were isolated from mutant mice on both BL6 and JAX backgrounds as well as corresponding wild-type mice, and the carotid sinus nerve activity was recorded *ex vivo*. CBs were challenged with graded hypoxia. Examples illustrating the effect of hypoxia (*p*O_2_ ∼40 mmHg) in Olfr 78 (BL6 and JAX backgrounds) and the corresponding wild-type mice are shown in Fig. 6 A-B and average data of CB sensory responses to graded hypoxia are presented in Fig. 6 C-D. The magnitude of sensory nerve response at given *p*O_2_ was significantly less in Olfr78 null CBs on both BL6 and JAX backgrounds as compared to corresponding wild-type CBs (Fig. 6 C-D).

**Fig. 6.**
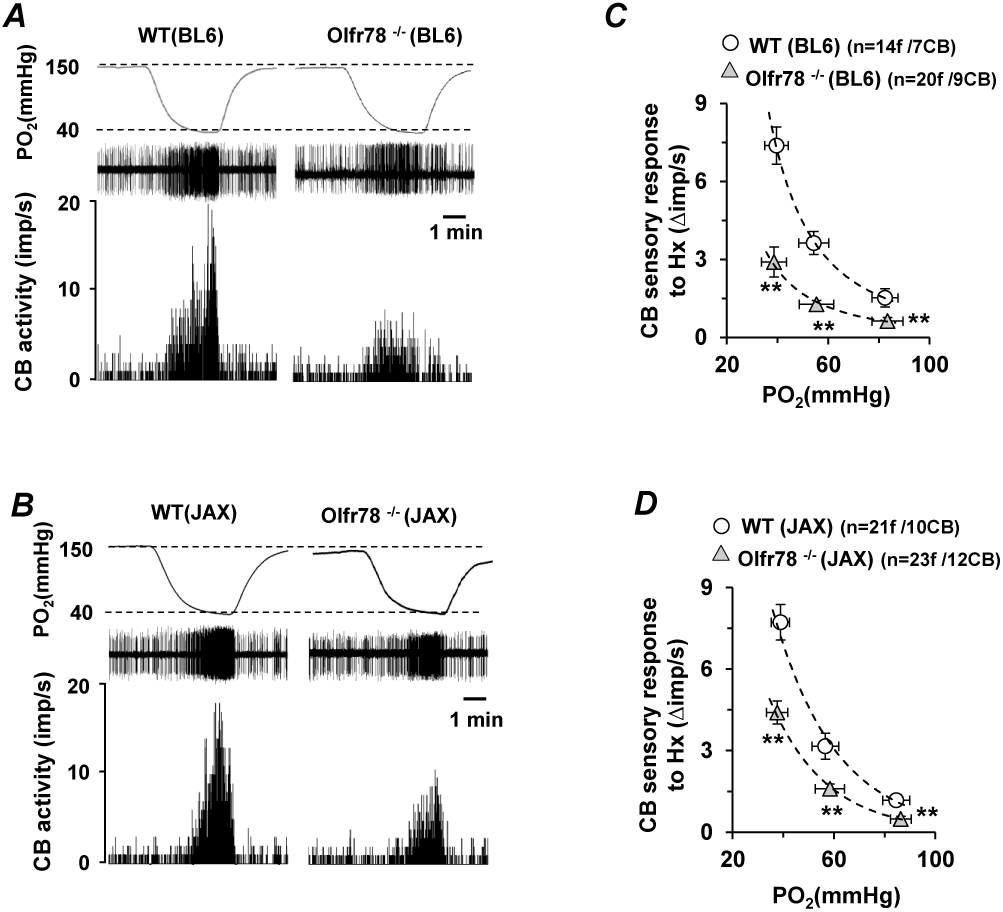
Carotid body (CB) sensory nerve response to hypoxia in wild type (WT) and Olfr78 null (Olfr78 ^-/-^) mice on BL6 (***A***) and JAX (***B***) background. *p*O_2_ (mmHg) in the superfusion medium measured by an O_2_ electrode placed closed to the CB (*top panels*), raw action potentials (*middle panels*) and integrated action potential frequency impulses per second (*bottom panels*). ***C-D***. Average data (mean ± SEM) of CB sensory responses to graded hypoxia presented as stimulus-evoked response minus baseline levels (Δ impulse/sec). Numbers in parenthesis represent the number of fibers and number of CBs. WT vs Olfr78 ^-/-^; Two-way ANOVA with repeated measures followed by Tukey test, ** P<0.01.

### [Ca^2+^]_i_ responses of glomus cells to hypoxia

We next determined whether Olfr78 is required for [Ca^2+^]_i_ response of glomus cells to hypoxia. Studies were conducted on glomus cells isolated from wild type and Olfr78 mutant mice on BL6 and JAX backgrounds, and were maintained overnight (∼18 h) in a growth medium. For these experiments, we chose *p*O_2_∼ 30 mmHg because severe hypoxia (*p*O_2_ < 30 mmHg) produces markedly reduced sensory nerve excitation(Peng et al. 2019).

Hypoxia increased [Ca^2+^]_i_ in wild-type glomus cells and this effect was attenuated in Olfr78 mutant mice raised either on BL6 or JAX background (Fig. 7A-D). In contrast, [Ca^2+^]_i_ elevation produced by 40mM KCl, a non-selective depolarizing stimulus were comparable between wild-type and Olfr78 null glomus cells (Fig. 7 A-D).

**Fig. 7.**
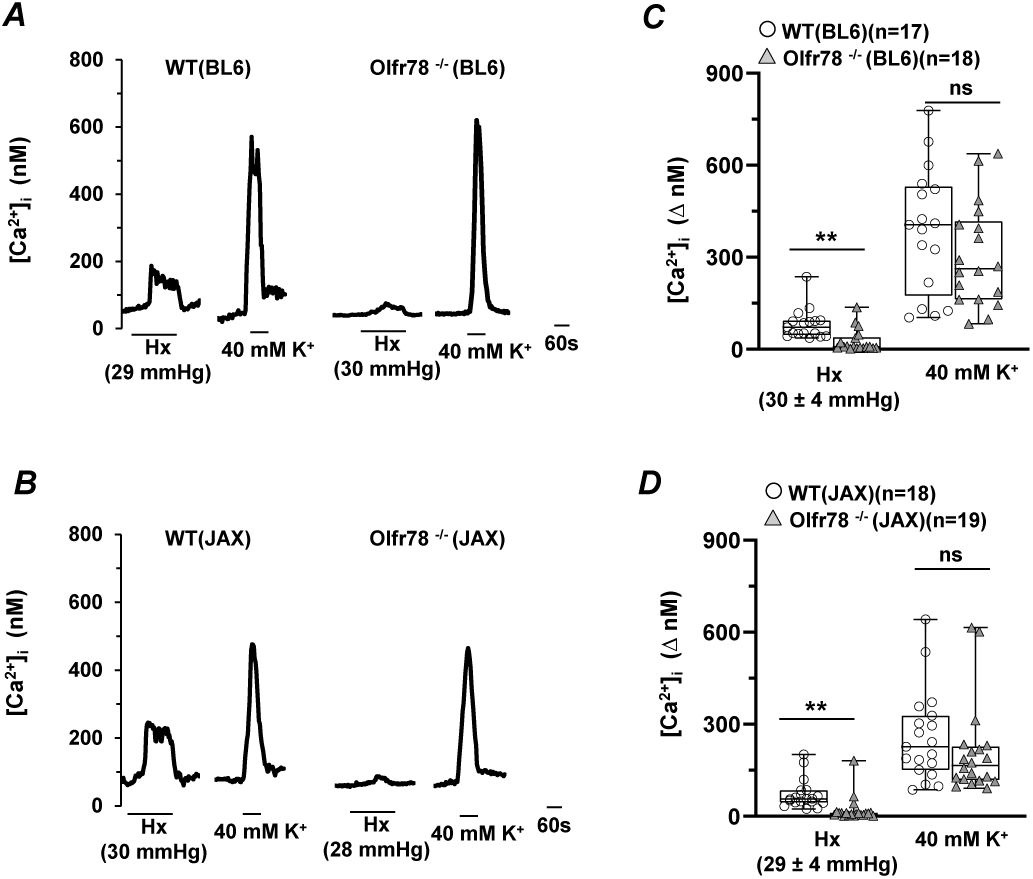
[Ca^2+^]_i_ responses of glomus cells from wild type and Olfr78 null mice on BL6 (***A***) and JAX (***B***) backgrounds to hypoxia (Hx) and 40mM KCl (K^+^). *p*O_2_ in mmHg represents partial pressure of O_2_ in the medium irrigating the glomus cells. ***C-D***. Box-Whisker plots with individual data points showing data of [Ca^2+^]_i_ responses of glomus cells to hypoxia and 40mM K^+^ presented as stimulus-evoked response minus baseline levels (Δ[Ca^2+^]_i_, nM). Number of glomus cells is presented in the parenthesis. WT vs Olfr78 null glomus cells, Mann-Whitney Rank Sum test, ** P<0.01, n.s. not significant, P>0.05.

## DISCUSSION

Consistent with the findings of Chang et al (Chang et al. 2015), we also found attenuated HVR, a hallmark of the CB chemo reflex, in un-sedated Olfr78 null mice as measured by whole body plethysmography. The reduced HVR was primarily due to attenuated respiratory rate rather than tidal volume and is reflected in minute ventilation which was normalized to body weight as well as oxygen consumption (VO_2_), the later a measure of convective requirement. Such an attenuated HVR was evident in Olfr78 null mice both on BL6 and JAX background strains (Table 1), thereby negating any potential confounding influence of background strains. These observations differ from those reported by (Torres-Torrelo et al. 2018), who monitored integrated respiratory rate as a measure of breathing in un-sedated mice. However, we found it is technically not possible to integrate respiratory rate in un-sedated mice, because of highly unstable breathing arising from sniffing and exploratory behavior, which was also reported by other investigators (Tankersley et al. 1994, Dauger et al. 1998, Tankersley et al. 1993). Because of the unstable breathing, it was necessary to choose segments of 15-20 consecutive breaths of stable breathing every minute for analyzing breathing during baseline and during hypoxia in un-sedated mice.

One might argue that our analysis of HVR is biased because of the choice of stable breaths. To circumvent this limitation, we further analyzed HVR by monitoring efferent phrenic nerve activity as an index of breathing in anesthetized mice. Our results showed a clear attenuation of HVR in anesthetized Olfr 78 null as compared to wild-type mice. Attenuated HVR in anesthetized Olfr 78 null mice is due to both reduced tidal volume as well as the respiratory rate as opposed to attenuated respiratory rate in un-sedated mice. Remarkably, stimulation of breathing by CO_2_ was unaffected in Olfr78 null mice measured either by plethysmography or by efferent nerve activity. Taken together, these results demonstrate selective impairment of HVR, which is a hallmark of the CB chemo reflex, in mice lacking Olfr78.

It is well established that depression of breathing evoked by brief hyperoxia is an indirect measure of peripheral chemo receptor sensitivity to oxygen both in humans (Dejours 1962), and in experimental animals (Kline et al. 1998). Olfr 78 null mice (both on BL6 and JAX background) displayed remarkable attenuation of breathing depression by hyperoxia, suggesting impaired peripheral chemo receptor, possibly CB sensitivity to O_2_. To further establish the effects of Olfr78 deficiency on CB function, we directly monitored the CB sensory nerve responses to graded hypoxia. It is clear from our results, CB sensory nerve response to any given *p*O_2_ is markedly attenuated in Olfr78 generated both on BL6 or JAX backgrounds. Consistent with the response of the CB sensory nerve activity, hypoxia-evoked [Ca^2+^]_i_ elevation of glomus cells is also impaired in Olfr78 null mice, whereas [Ca^2+^]_i_ elevation evoked by KCl, a non-selective depolarizing stimulus was unaffected. These findings are consistent with the earlier observations by Chang et al. (Chang et al. 2015), and demonstrate that Olfr78 is an important component of the CB hypoxic sensing pathway.

In striking contrast to present observations, Torres-Torrelo et al found that hypoxia-evoked [Ca^2+^]_i_ elevation was unaffected in Olfr78 null glomus cells (Torres-Torrelo et al. 2018). A major difference between our study and by Torres-Torrelo et al (Torres-Torrelo et al. 2018) is the severity of hypoxia used for evoking [Ca^2+^]_i_ elevation in glomus cells. Torres-Torrelo et al used 10-15 mmHg pO_2_(Torres-Torrelo et al. 2018), whereas we employed *p*O2 of ∼30 mmHg. We recently reported that severe hypoxia (i.e., below pO2 of 30mmHg) although evokes [Ca^2+^]_i_ elevation in glomus cells but produces only a weak stimulation of the CB sensory nerve activity and depress breathing rather than stimulation (Peng et al. 2019). Therefore, we employed a medium pO_2_ of ∼30 mmHg for eliciting Ca^2+^]_I_ response of glomus cells, which produces robust CB sensory nerve excitation (Fig. 6), which is an essential pre-requisite for triggering reflex stimulation of breathing. It follows that Olfr78 play an important role in the CB excitation to a physiologically relevant hypoxia but not to near anoxic stimulus.

## Acknowledgement

This work was supported by National Institutes of Health grants P01-HL-44454.

